# Exposure to a low dose mixture of endocrine disrupting chemicals alters the brain transcriptome and animal behavior

**DOI:** 10.64898/2026.02.10.705055

**Authors:** Alekh Paranjapye, Rili Ahmad, Camille N. Quaye, Andre L.G. Rico, Nicole Palmiero, Rebecca A. Simons, Yu-Chin Lien, Molly Hall, Erica Korb

**Affiliations:** Department of Genetics, University of Pennsylvania Perelman School of Medicine, Philadelphia, PA, USA; Epigenetics Institute, University of Pennsylvania Perelman School of Medicine, Philadelphia, PA, USA; Centre for Women’s Health and Reproductive Medicine, University of Pennsylvania Perelman School of Medicine, Philadelphia, PA, USA; Department of Pediatrics, University of Pennsylvania Perelman School of Medicine, Philadelphia, PA, USA

**Keywords:** Endocrine disrupting chemicals, mouse neuron culture, mouse development, behavior, single nuclei RNA sequencing, cell-cell signaling, sex differences

## Abstract

Exposures to pervasive chemical toxicants such as endocrine disrupting chemicals (EDCs) are associated with adverse neurological and neurodevelopmental deficits. Although EDCs are widespread as sparse mixtures in the environment, most research has focused on single chemicals at high concentrations. Here, we studied the effects of ldEDC: a low-dose mixture of widely prevalent toxicants at doses representative of normal human exposure levels. Primary cultured mouse neurons treated with ldEDC exhibited altered gene expression compared to vehicle controls in genes critical for neuron activity, indicating low doses EDCs can affect neuronal function directly. We next tested persistent exposure through the maternal diet to define perinatal effects on offspring. Exposed offspring exhibited differences in development, tactile sensitivity, and sex-specific changes in motor behavior. Cortical single-nuclei sequencing identified broad transcriptomic changes, particularly in distinct cortical layer subpopulations, excitatory neurons, and astrocytes. Cell-cell signaling between neurons and non-neuronal populations were altered in exposed mice, specifically in pathways associated with cellular adhesion. Transcriptomic differences were also sex-specific. Together, these *in vitro* and *in vivo* findings reveal molecular and phenotypic consequences of EDC exposure at a mixture of doses well below commonly studied levels and highlights common functional pathways of susceptibility.

## Introduction

Exposures to endocrine-disrupting chemicals (EDCs) and environmental pollutants are associated with increased risk for neurodevelopmental disorders (NDDs)(1–7). Perinatal and early life exposure to EDCs are linked to impaired cognitive functioning and aberrant behaviors throughout adolescence(8,9) and adulthood(10–12). Correlations have been shown between EDC burden and the severity of disorders such as attention deficit hyperactivity disorder (ADHD)(13–15) as well as intellectual disability(16). Although constrained and not necessarily causative in isolation, these findings suggest that EDCs can act as risk factors due to their potential to negatively affect brain development and health, especially if paired with causative genetic factors. Despite the growing evidence for connections between NDDs and toxicants such as bisphenols(17) such as bisphenol A (BPA)(18), phthalates such as DEHP(19), and perfluoroalkyl substances such as PFOS(13,20–22) that can act as EDCs and through other mechanisms, their effects on neuronal function are not fully understood. Even less well understood are their effects at the low levels found in humans and in the environment.

Most studies of EDCs examine the consequences of single chemicals and may not capture effects of exposure to multiple chemicals commonly occurring in the environment that cannot be predicted by any individual component alone. Thus, while studying single compound exposures is critical for defining mechanisms of action, EDC mixtures more closely represent normal human exposures and are of particular relevance to public health(23–27). Furthermore, mixtures of EDCs at environmental levels below individual estimated ‘no observed adverse effect levels’ (NOAELs) may have determinantal impacts only observable in the combined dosage(28). In support of this, perinatal exposure to a representative EDC mixture termed NeuroMix caused physiological and behavioral changes in rats(29). This work found that even at low dose of the mixture and a short window of exposure, pups exhibited significant differences in development, behavior, and the brain transcriptome.

We sought to expand on this landmark work by utilizing a modified version of NeuroMix termed ldEDC, composed of a subset of NeuroMix toxicants of current high public health concern and known associations with NDD phenotypes(20,19,12,18,7). Further, we transitioned to mice to allow for future studies pairing exposure with genetic manipulations that are causative for NDDs. First, we tested the effects of exposure on mouse primary cultured neurons in isolation to define the direct effects of ldEDC on neurons. We then performed perinatal treatment using a passive oral route through the diet to more accurately replicate normal human exposures. We tested offspring growth, developmental milestones, and behavior. Additionally, we performed a highly multiplexed single-nuclei sequencing strategy SPLiT-seq to profile the cortical transcriptome of exposed and non-exposed mice to define the molecular consequences of exposure. Our findings show that even at doses well below the individual NOAEL of each component, low-dose mixtures of pervasive EDCs cause subtle but broad and sex-specific changes in gene expression. Disruptions were found in both cultured neurons and in the brain and linked to network disruptions. Further, we detected changes in growth, developmental milestones, and movement, demonstrating functional consequences of exposures. Together, these results indicate that exposure to EDCs in mice under the NOAELs can lead to phenotypic consequences when occurring in combination.

## Results

### Transcriptomic analysis of cultured neurons exposed to low-dose endocrine-disrupting chemical mixture ldEDC

Previous work using NeuroMix found that the perinatal exposure to the low-dose EDC mixture induced anxiety and social phenotypes, which were associated with changes in gene expression in the medial amygdala(29). To expand on these results in mice, we first created a version of the mixture called ldEDC (**Supplemental Data Table 1**). ldEDC contains a subset of chemicals used in NeuroMix, including two bisphenols (BPA and BPS), two phthalates (DEHP and DBP), a diphenyl ether (PBDE-47), and a polyfluoroalkyl substance (PFOS). These chemicals were selected as they are all pervasive in the environment(4,30,31), still in circulation across industrial nations including the United States, are detectable in humans, and are associated with neurobiological deficits in mammals(32–34,29). We first tested exposure x exposure interactions were computationally validated using the Integrative Genome-Exposome Method (IGEM) system, which leverages exposomic interactions from multiple public databases(35) and observed reported interactions between all six EDCs and at least one other EDC tested (**Fig. 1A**). Next, we examined whether these toxicants may have overlapping functional effects on by examining reported gene interactions for each EDC in the Comparative Toxicogenomics Database(36). We found overlapping gene interactions between all EDC combinations with over 800 genes interacting with all six chemicals (**Fig. 1B**).

**Figure 1:**
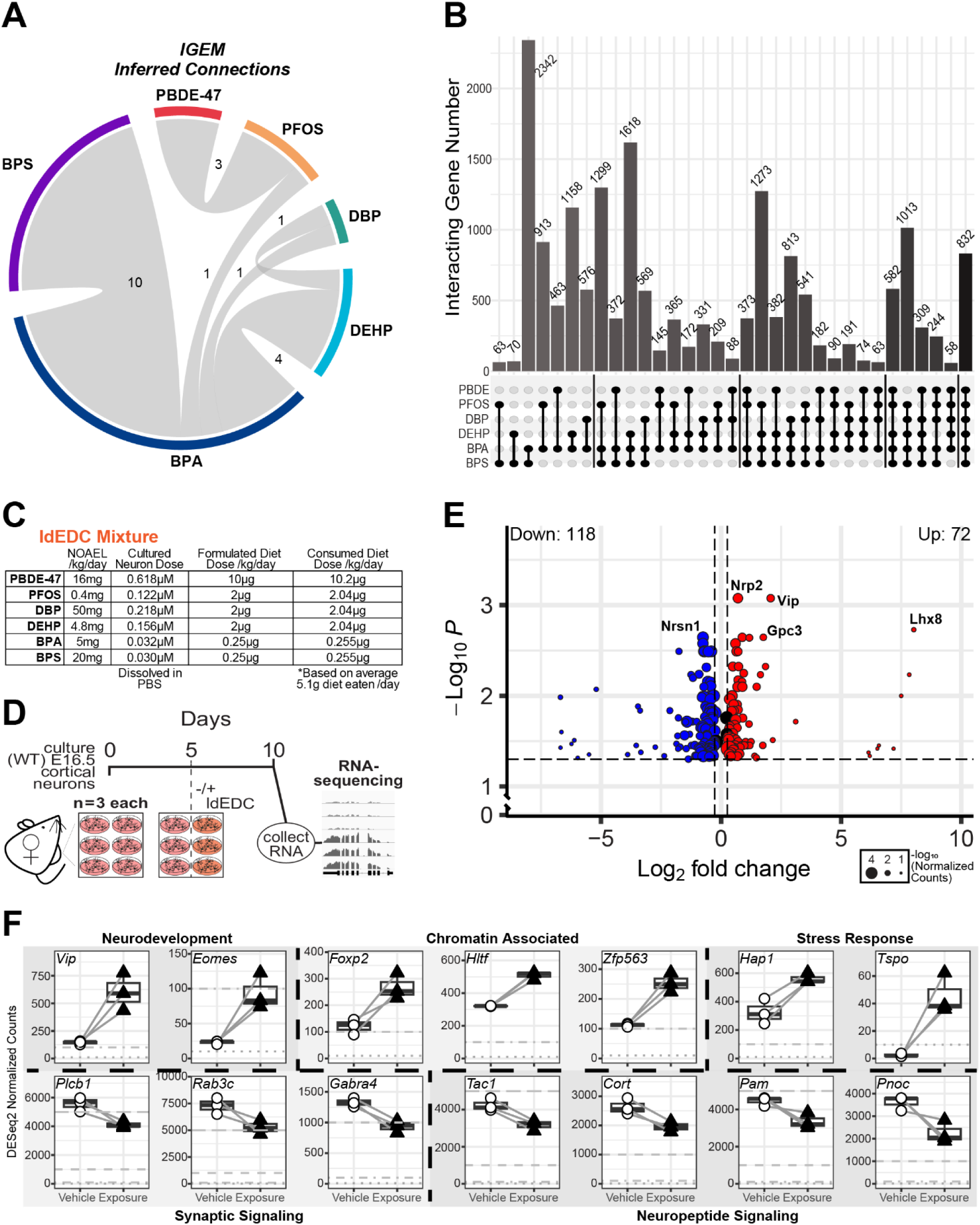
Gene expression analysis of neurons exposed to a low dose endocrine disrupting chemical mixture. **A.** Exposure x Exposure interactions identified using the Integrative Genome-Exposome Method (IGEM) on the CTD. Only those interactions with curated, pre-processed vocabulary and normalized to selected internal concepts were selected. Repeated co-occurrences were counted once per record. **B.** Upset plot of the overlap of genes with characterized interactions in the Comparative Toxicogenomics Database (CTD) with each of six common endocrine disrupting chemicals (EDCs). **C.** Table detailing the no-observed-adverse-effect level, *in vitro* concentration, or *in vivo* consumed weight of each component of the ldEDC mixture in this study. **C.** Schematic of the experimental timeline for the comparison of transcriptomes between neurons treated with vehicle or the EDC mixture ldEDC. (n = 3) biological replicates (neuronal cultures derived from different liters). **D.** Normalized counts of select DEGs in the neuronal exposure, organized by functional categories. Paired replicates are connected between conditions. **E.** Volcano plot of differentially expressed genes (DEGs) following ldEDC exposure (|fold change| ≥ 1.2, adj(p) ≤ 0.05).

To first determine if this truncated mixture affects the biology of neurons, we used primary neuronal cultures derived from embryonic day 16.5 mouse cortical tissues to generate a highly pure neuron population and treated neurons with vehicle or a solution of ldEDC (**Fig. 1C-D**). This system allows us to test exposure within a pure neuronal population without the complexity and heterogeneity of *in vivo* exposures, in genetically identical neurons derived from the same litters, and without confounding effects of mixed backgrounds from patient samples. It also provides a highly controlled system with control and treated neurons derived from the same embryo and removes litter-to-litter variability inherent in perinatal exposure paradigms. Further, neuron cultures are free of hormonal effects that may be caused by birth, the early days of life, and adolescence which could indirectly affect key pathways that could be disrupted by EDCs. Finally, treatment of primary culture populations after neuronal identity is established and cells are fully post-mitotic allows for specific examination of effects on neuronal maturation in the absence of confounding effects on neuronal identity.

We cultured neurons from wildtype cortices (3 biological replicates composed of 4 mixed cortices each) and treated them with either vehicle control or a solution of ldEDC for 5 days in vitro (DIV). We then maintained the culture for an additional 5 days before collecting RNA and performing bulk RNA-seq. A moderate number of genes were differentially expressed, with a greater number downregulated than upregulated (**Fig. 1E, Supp Fig. 1A**). Although no gene ontology terms were enriched amongst significantly up or downregulated genes, there was marked differential expression of several genes encoding proteins essential in important neuronal functions (**Fig. 1F, Supp Fig. 1B**). These include regulators of neuronal fate such as *Eomes* and *Foxp2,* transcription factors such as *Foxp2*, synaptic signaling components like *Rab3c* and *Gabra4*, neuropeptide precursors such as *Tac1*, and G-protein regulators *Rgs2* and *Rgs13*.

### Perinatal ldEDC exposure has subtle effects on mouse development and behavior

We next moved into an in vivo system to determine the consequences of persistent exposure to EDCs in mice. To accurately reflect oral ingestion of low-levels of EDCs over long periods of time, we generated a custom diet of low-phytoestrogen feed with or without ldEDC (**Supplemental Data Table 1**). We first tested exposure in pregnant female mice in a preliminary cohort on a diet of 1X of each compound in the mixture to determine health effects. We also tested a 0.1X mixture containing one-tenth of each chemical in the formulation to model regular low-dose exposures that may be more typical of regular human exposures. Female mice receiving a 1X dosage exhibited substantial weight loss, stunting, and incapability to become pregnant and were sacrificed (**Supp Fig. 2A**). T Conversely, the 0.1X dosage, showed limited effects in adult mice and was selected for subsequent testing. Female mice were placed on the diet two weeks prior to mating, during pregnancy, and through weaning (three weeks after delivering pups) (**Fig. 2A**). This timeline allowed for exposure across development and early life of the F1, emulating low dose but consistent EDC exposure.

**Figure 2:**
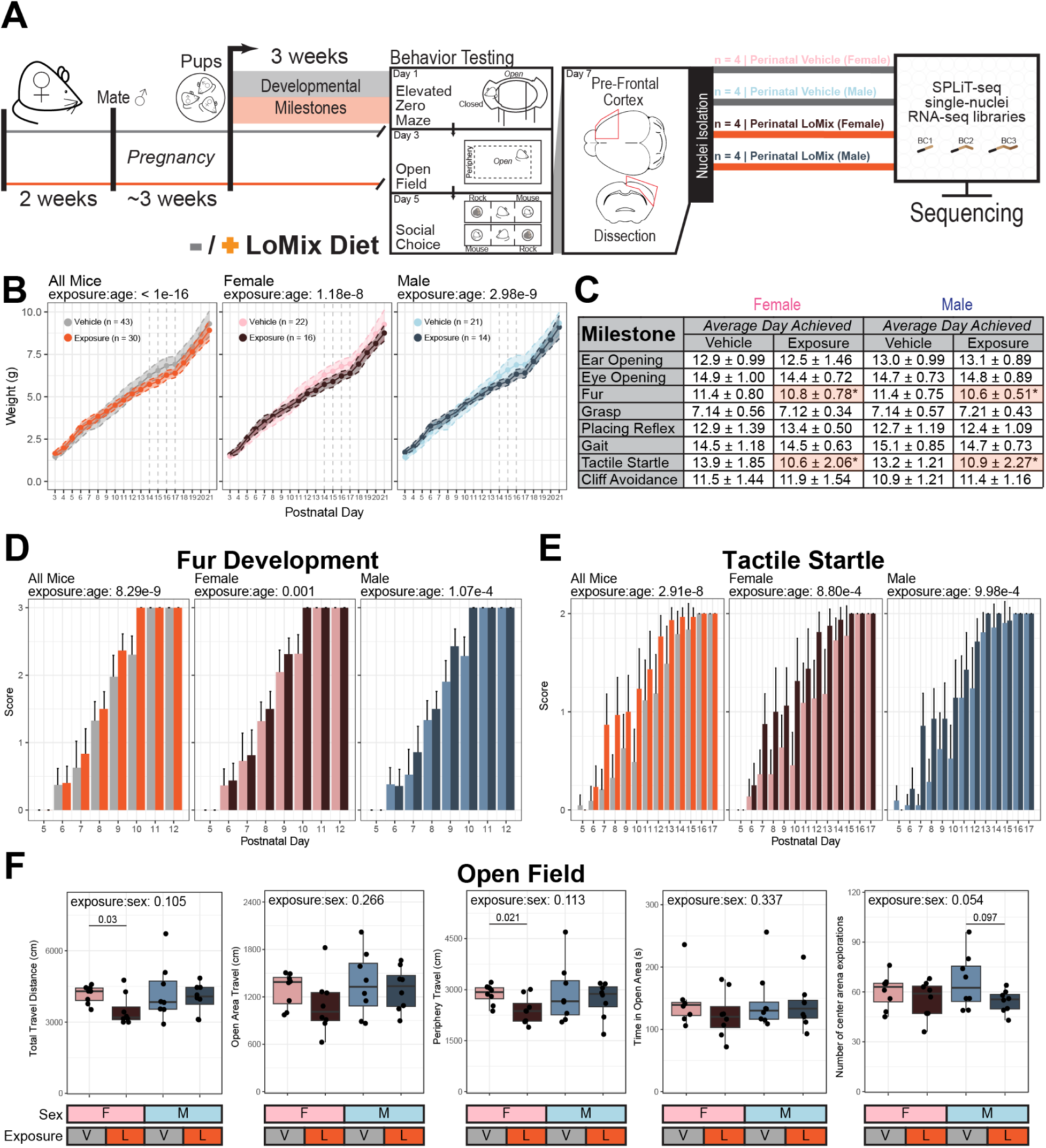
Developmental and behavioral differences in perinatally ldEDC exposed mice. **A**. Schematic of the experimental timeline for the ldEDC diet administration, developmental and behavior testing, and single-nuclei RNAseq. **B**. Weight during first three weeks post-birth in male and female mice (repeated measures ANOVA above, dashed lines indicate unpaired two-tailed t-test < 0.05) (**C**. Developmental milestone achievements in female and male mice (female[vehicle: n = 22, exposure: n = 16]; male[vehicle: n = 21, exposure: n = 14] unpaired two-tailed t-test). **D**. Fur development score at each tested day after birth (scores: 0 = no fur, 1 = dorsal pigment appearance, 2 = dorsal and ventral pigment appearance with short dorsal hairs, 3 = fur appearance). **E**. Tactile startle score at each testing day after birth (scores: 0 = no response to air puff, 1 = slight motion response, 2 = exaggerated jumping response). **F**. Measures from the open field test (repeated measures ANOVA above each, Kruskal-Wallis Test between pairs).

All F1 offspring (5 cohorts vehicle [22 females and 21 males], 5 cohorts exposure [16 females and 14 males]) were weighed every day (**Fig. 2B**) and developmental milestones were tracked (**Fig. 2C, Supp Fig. 3A-B**) as detailed in Heyser (2004)(37). Both female and male pups gained weight at slower rates than those with the vehicle diet, although this deficit was lost between days 17 and 18 after birth which roughly corresponded to when the pups were able to eat solid food. ldEDC-exposed pups did not have any deficits in most milestone achievements including physical development (ear and eye opening), motor activity (gait), some reflexes (grasp, placing), or proprioception (cliff avoidance, righting, negative geotaxis). However, mice exposed to the EDCs developed fur faster than control counterparts (**Fig. 2D**) and exhibited a pronounced startle response at an earlier age (**Fig. 2E**). This suggests that EDC exposure can cause disruptions to normal developmental trajectories and behavioral responses.

We next performed a series of behavioral tests to determine if perinatal EDC exposure results in differences in anxiety or social phenotypes, representative of neurodevelopmental impairment. There were no differences observed between groups in the elevated zero maze (**Supp Fig. 3C**) or social choice tests (**Supp Fig. 3D**). In an open field experiment, there were no significant differences in the open area time across paradigms. However, females exposed to ldEDC travelled less than controls, primarily around the periphery (**Fig. 2F**). We also observed a nonsignificant trend toward a sex-exposure interaction in the number of times entering center of the area by males. This suggests that while mice are generally healthy and show no major anxiety or sociability deficits, even very low doses of EDCs can affect developmental behaviors and have sex-specific effects on motor behavior.

### Single-nuclei RNA-seq reveals cell-specific consequences of perinatal ldEDC exposure

Exposure to EDCs, including in mixtures, is known to cause epigenetic and transcriptional changes that persist through early life(32,38). To characterize these changes in the ldEDC exposed mice, we performed single-nuclei RNA sequencing (snRNA-seq) of the prefrontal cortices of our F1 offspring after the behavior circuit (**Fig. 2A**). Due to known sex differences with toxicant exposure(39–42), we chose 4 male and 4 female mice from each condition based on the behavior data, selecting mice to avoid outliers in open field and social choice experiments. To minimize batch effects across multiple chips in traditional droplet-based assays, we processed the isolated cortical nuclei in parallel using SPLiT-seq with the Evercode WT v3 kit (Parse Biosciences). We identified clusters broadly capturing 4 canonical inhibitory neuron types (*Gad1/2*+), 2 glycine-GABA inhibitory neuron types, 6 excitatory neuron types representing those of 5 cortical layers (*Slc17a7*+), 2 oligodendrocyte populations (*Mag*+), a cluster representing astrocytes and radial glia cells (*Atp1a2*+), microglia (*Inpp5d*+), and endothelial cells (*Fzd3*+) (**Fig. 3A, Supp Fig. 4A**).

**Figure 3:**
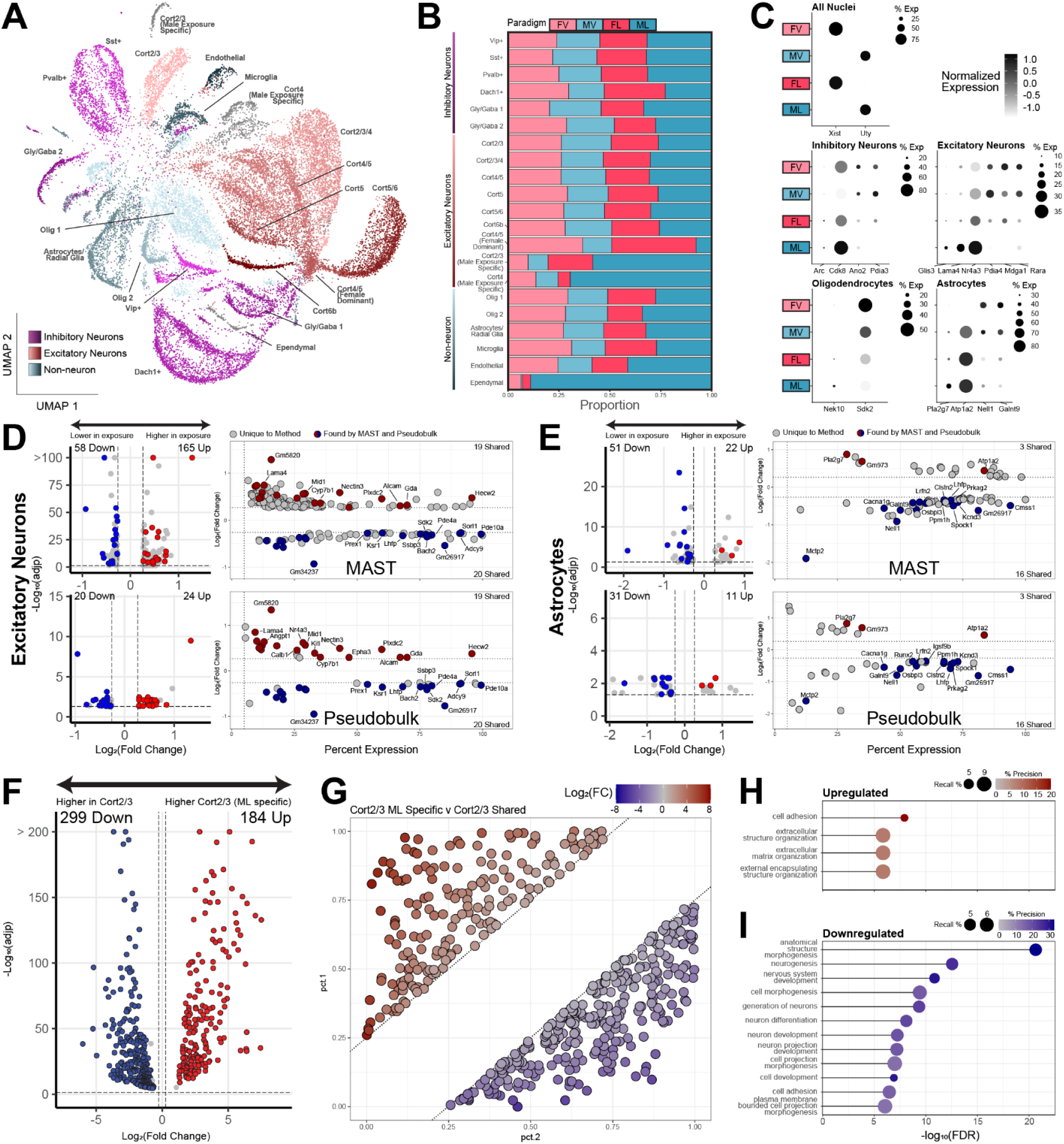
Cortical single nuclei transcriptomic responses to exposure. **A**. UMAP of single-nucleus transcriptomic profiles from female or female, vehicle or ldEDC exposed mouse cortices (n = 4 for each sex-condition combination). **B**. Stacked bar chart of the proportion of each sex-condition combination for each cluster (FV: female-vehicle, FL: female-ldEDC, MV: male-vehicle, ML: male-ldEDC). **C**. Dotplots of normalized expression and percent expression per condition for select differentially expressed genes. **D-E**. Volcano and scatter plots of differentially expressed genes between exposure and vehicle for all excitatory neurons (**D**) or astrocytes (**E**). Points are colored if the gene was significant (|fold change| ≥ 1.2, adj(p) ≤ 0.05) in both the MAST and pseudobulk analysis. **F**. Volcano plot of genes differentially expressed in the ML-specific cortical 2/3 excitatory neuron cluster compared to the condition-balanced cortical 2/3 cluster. **G**. Scatterplot of genes by their percent expression in the ML-specific cortical 2/3 cluster (pct.1) or condition-balanced cortical 2/3 cluster (pct.2). Only those genes with an |fold change| ≥ 1.2, adj(p) ≤ 0.05, and minimum pct = 0.25 are shown. **H-I**. Gene ontology analysis of significantly up (**H**) or down (**I**) differentially expressed genes between the cortical 2/3 clusters. Recall is the proportion of functionally annotated genes in the query over the number of genes in the GO term. Precision is the number of genes found in the GO term over the total number of genes in the query.

Interestingly, we also identified 3 clusters that were primarily enriched for nuclei from perinatally exposed male mice (**Fig. 3B, Supp Fig. 5B**). Two of these clusters were excitatory neurons corresponding to cortical layers 2/3 and 4, and the third cluster was most closely classified as a population of brain ependymal cells although this cluster had too few nuclei for any analysis and was omitted from downstream processing. Additionally, we identified a subcluster of the cortical 4/5 excitatory neurons enriched for nuclei from female samples from both conditions. No other clusters were significantly overrepresented by any single condition (**Supp Fig. 4C**). These findings suggest that low-dose EDC exposure is sufficient to cause subclustering of specific susceptible populations in a sex-specific manner.

We first performed pairwise comparisons between vehicle and exposure nuclei separated by populations encompassing all cells of the same type (**Fig. 3C-E**) and excitatory neuron clusters (**Supp Fig. 4D**). To minimize false positive DEGs often found by the default methods of differential gene expression analysis, we compared conditions using both MAST and pseudobulk, and highlighted transcripts identified by both. A subset of DEGs were found up- or downregulated in more than one excitatory cell cluster (**Supp Fig. 5E-F**). Fewer DEGs were identified for inhibitory neurons (**Supp Fig. 5A**), combined or separated by subcluster.

To determine what transcriptomic features differentiate the male-exposure-specific clusters, we performed pairwise comparisons between these clusters and the cortical layer clusters of closest identity (**Fig. 3F-G, Supp Fig. 5B-C**). The male-exposure-specific cortical 2/3 cluster was most substantially different from the shared cortical 2/3 layer with over 184 and 299 up and downregulated DEGs in comparison, respectively. Upregulated genes were enriched for those that encode extracellular membrane components (**Fig. 3H**) while the unique cluster exhibited substantial downregulation of genes encoding proteins involved in neuronal development, structure, and projection (**Fig. 3I**). The male-exposure-specific cortical layer 4 cluster had fewer DEGs but, interestingly, were enriched for neuronal signaling components (**Supp Fig. 5D**).

### ldEDC exposure alters cortical cell-to-cell signaling pathways

Because we identified differential gene expression of transmembrane-encoding proteins in pairwise comparisons, we analyzed differences in receptor-ligand signaling pathways using CellChat(43). We did not find differences in the total number of inferred signaling interactions nor cumulative interaction strength between vehicle and exposure. However, numerous connections between clusters were altered with exposure causing both increases and decreases in the number and strength of specific cluster interactions (**Fig. 4A-B**). Exposure resulted in increases in signaling from inhibitory neurons across cell populations, as well as a parallel reduction in signaling from cortical layer 5 and most glial cells. Next, we examined the differential signaling pathways and found only slight differences between conditions across numerous classes of receptor-ligand groups (**Supp Fig. 6A**). We quantified the cumulative information flow from all pathways contributing at least 25% of the total signaling weight and found several with significantly greater and lower signaling between conditions (**Fig. 4C, Supp Fig. 6B**). The most downregulated pathways following exposure were adhesion associated, such as adhesion G signaling (ADGRG), semaphorin (SEMA4), and tenascin (**Fig. 4D, Supp Fig. 6C**). Among those with increased signaling were Platelet-derived Growth Factor signaling (PDGF), growth arrest specific signaling (GAS), neurotrophin (NT) and cholesterol (**Fig. 4E, Supp Fig. 6D**).

**Figure 4:**
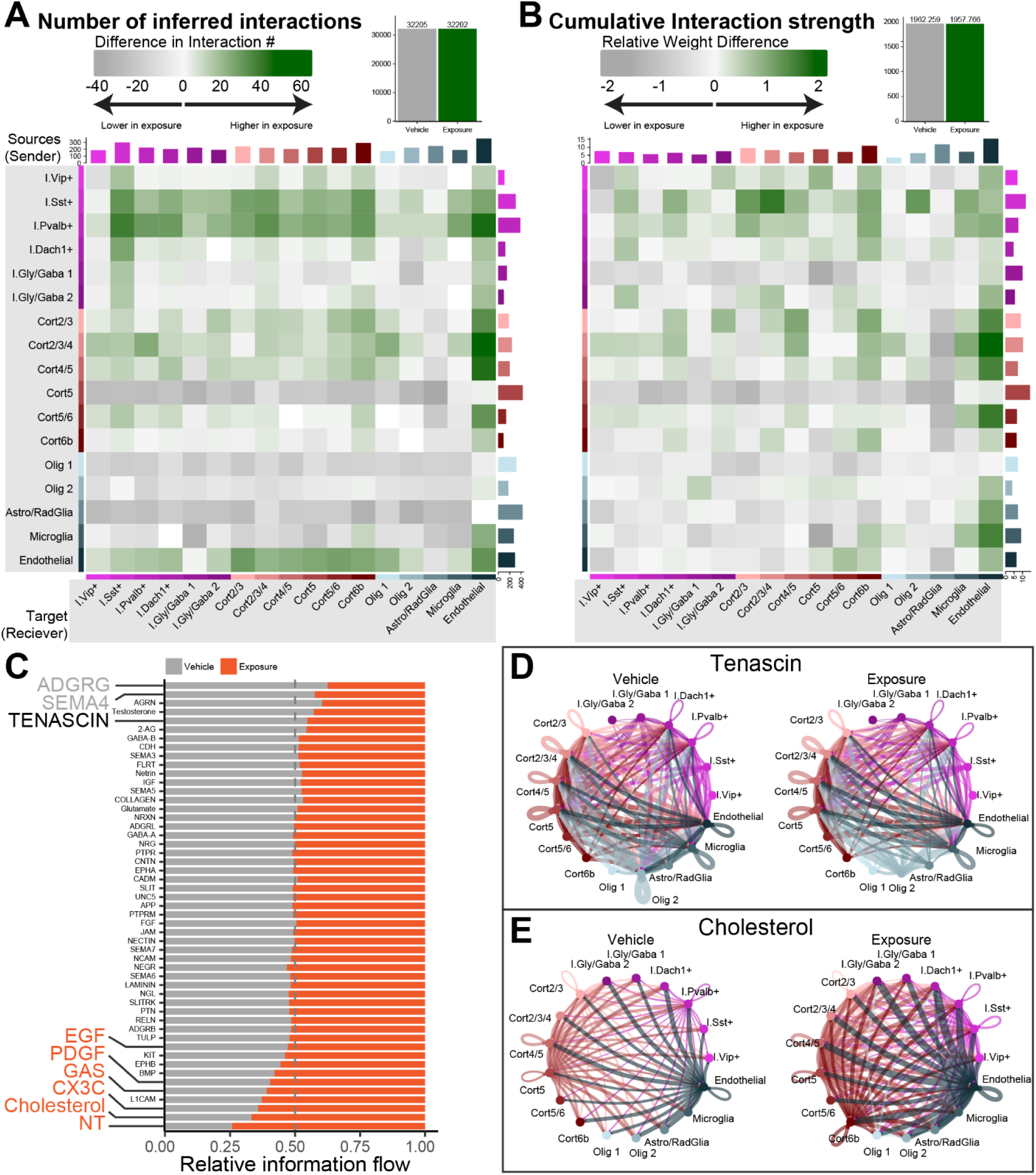
Cell-cell communication changes between vehicle and exposure cortices. **A**. Total number of significant signaling interactions identified between vehicle and exposure nuclei, and heatmap of differences between conditions observed from a ligand-bearing source (row cluster) to receptor-bearing target (column cluster). The absolute value of the sum of differences for each source or target is shown in the horizontal or vertical bar plot next to heatmap, respectively. **B**. Relative weight total and differences between source and target clusters as calculated by law of mass communication probability calculations. **C**. Stacked bar plot of the relative information flow from signaling pathways contributing to ≥ 25% of the total signaling weight found in vehicle or exposure conditions. Colored and expanded names indicate pathways that are differentially enriched in vehicle (grey) or ldEDC-exposed mice (green). **D-E**. Chord diagrams illustrating significant interactions of the tenascin (**D**) or cholesterol (**E**) signaling pathways between clusters. Colors indicate the source of the L-R pair and the size of each chord indicates the cumulative weight.

### ldEDC transcriptomic signatures are unique between males and females

Due to the known sex differences in EDC effects and those we observed in this study, we stratified the single-nuclei dataset by sex and performed the same stringent pairwise comparisons to determine sex-specific DEGs by cell type (**Fig. 5**). We found several more genes differentially expressed in inhibitory neurons in females compared to males. (**Fig. 5A-B**). As in the combined sex analysis, the greatest number of DEGs overall were found between vehicle and exposure in excitatory neurons (**Fig. 5C-D**) and we again detected numerous genes primarily downregulated in astrocytes (**Fig. 5E-F**). Additionally, when separated by sex, we identified female and male-specific DEGs in oligodendrocytes, none of which were detected when sexes were combined (**Fig. 5G-H**). Female-specific downregulated genes were the only groups with computed gene ontology enrichment, with genes encoding proteins associated with the dendritic spine (**Supp Fig. 7A**). We also performed STRING protein-protein interaction analysis of all DEG groups and found significant modules in the male-specific excitatory neurons, which included a class of integrins and endoplasmic reticulum chaperones (**Supp Fig. 7B-C**). Unexpectedly, we detected little direct overlap between DEGs in female or male backgrounds. Further, we found several DEGs significantly upregulated in one sex and downregulated in the other such as Cux1 in excitatory neurons (**Fig. 5E-F**).

**Figure 5:**
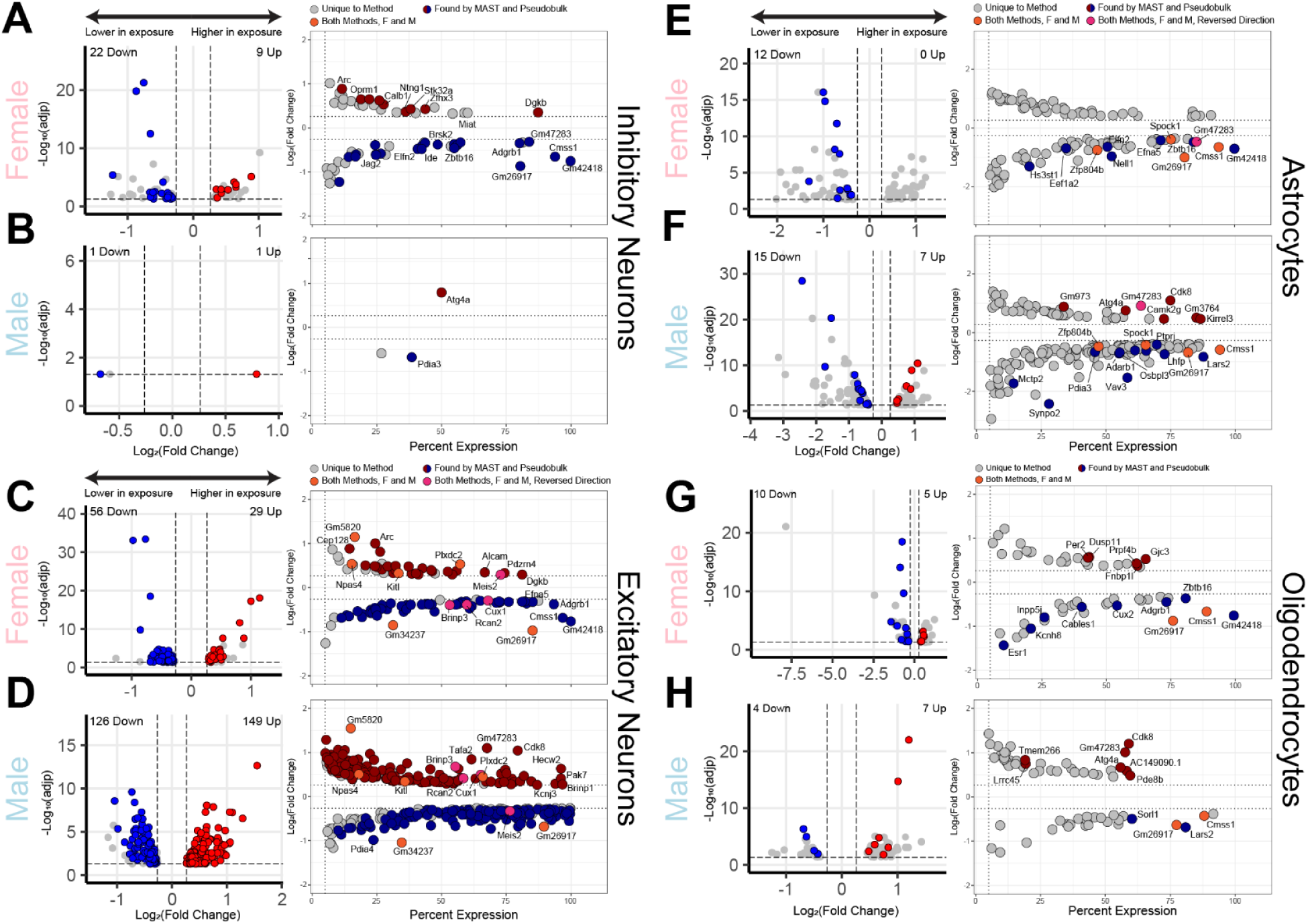
Sex-specific differential gene expression across brain cell types. **A-H.** Volcano and scatter plots of differentially expressed genes between exposure and vehicle for all inhibitory neurons (**A-B**), excitatory neurons (**C-D**), astrocytes (**E-F**), or oligodendrocytes (**G-H**) separated by only female or male nuclei. Points are colored if the gene was significant (|fold change| ≥ 1.2, adj(p) ≤ 0.05) in both the MAST and pseudobulk analysis. Orange points are genes found differentially expressed in both female and male conditions with the same directionality and purple points are found in both but opposite directionality.

We next sought to compare DEGs detected in young adult cortical tissue from single-nucleus RNA-sequencing with those identified in maturing neurons generated from E16.5 cultures. There was little overlap of DEGs identified from all excitatory neurons or sex-stratified sets with those found in cultured neurons (**Supp Fig. 7F-G**). Given the differences between systems, analysis methods, developmental timepoint, and processing ability of exogenous compounds in neurons grown in isolation vs whole animals, this is not unexpected. Together, these analyses define common and sex-specific gene expression changes in the brain following toxicant exposure. Further, these approaches identify sensitive functional groups and networks that indicate that even low-dose mixtures of compounds can elicit robust changes in gene expression in neurons.

## Discussion

Most studies examining the consequences of exposure to EDCs and other toxicants on neurons are limited by examining only single chemicals, and often at levels higher than those found in humans. This limits our understanding of the consequences of normal human exposures as multiple toxicants with similar effects at low doses may have effects that are not predictable from single chemical exposure. Here, we demonstrate the molecular effects of persistent exposure to a low-dose EDC mixture, (ldEDC), *in vitro* as well as phenotypic effects *in vivo*. In this study, we show that exposure had modest but widespread effects on gene expression in primary mouse neurons, specifically affecting key genes involved in neuronal development and the activity response. Using an *in vivo* system of developmental exposure to ldEDC, we detected differences in development and behavior with several sex-specific phenotypes. Single-nuclei RNA sequencing of the prefrontal cortices of these mice showed changes in gene expression in excitatory neurons as well as glial cells, with effects on genes relevant to neuronal networks and intracellular signaling pathways. We further detected substantial sex-specific changes in gene expression. These findings provide evidence that low doses of EDCs and other prevalent chemicals in combination can elicit changes that affect gene expression in neurons and behavioral outcomes in mice.

Similar to previous work studying the effects of a low-dose EDC mixture, we chose an oral route of perinatal exposure in our *in vivo* system to emulate common human exposures to toxicants. At a much lower dosage than other groups, we observed significant differences between perinatally exposed pups and their vehicle counterparts in the absence of changes in viability or major health deficits. Exposed pups exhibited a tactile startle response 1 to 2 days earlier than controls, which may be relevant to increased tactile sensitivity found in neurodivergent infants and children(44,45). Additionally, fur development occurred earlier which similarly could relate to the relationship between toxicant exposure and earlier onset of puberty(46–48). Also noteworthy is that the ‘1X’ dosage of EDC mixture used here, designed to be below NOAEL levels, was sufficiently toxic to preclude subsequent testing in mice.

To the best of our knowledge, this study provides the first high-throughput transcriptomic profiling of long-term, low-dose EDC exposure. Through single-nuclei RNA-seq, we observed minor changes in gene expression in glial clusters as well as excitatory neurons with distinct, male-exposure-specific signatures causing clusters to deviate from their closest populations. These findings point to lasting transcriptomic changes in neurons caused by low-dose exposure during development. Additionally, these changes extend to receptor-ligand pairs between neurons and non-neuronal cells, which mediate the important signaling functions of the brain. It is also worth noting, however, that many DEGs detected in this study may represent compensatory or protective mechanisms activated by neurons in response to chemical exposure and thus are not expected to be universally deleterious.

The broad but modest effects of low-dose exposures may not be sufficient to cause robust phenotypic consequences. However, they could indicate a predisposition for more severe phenotypes when combined with mutations in genes associated with neurodevelopmental disorders that have highly variable outcomes. Given that many NDDs such as ASD are predominantly caused by genetic risk factors, these findings could elucidate pathways of interaction with genetic factors associated with neurodevelopmental disorders. Other groups have examined interactions between EDC exposure and genetic perturbation in causing more severe phenotypes(49–52), but this has primarily been in high doses of single chemicals. While we found little direct overlap between DEGs and NDD-risk genes, we detected downregulation of several risk genes, including transcription factors such as Special AT-rich Sequence Binding Protein 1 and 2 (*Satb1/2*) and POU class 3 homeobox 3 (*Pou3f3*), as well as synaptic proteins like synaptojanin 1 (*Synj1*). Overall, our work highlights subtle but detectable effects of EDC mixtures in the environment that could contribute to detrimental effects, particularly when co-occurring with genetic risk factors. Future studies testing gene-environment interactions will be critical to identifying such molecular and behavioral consequences of these combinations.

When separating by sex, we were able to identify substantially more DEGs across neurons and glial cells as result of exposure. Unexpectedly, there was little overlap between the female and male-specific DEGs, indicating distinct molecular pathways affected by exposure in the brains of young mice. This corresponds to differences we and others have found at the phenotypic level between sexes and exposures(39,29,42). Interestingly, the only DEGs found in *in vitro* and *in vivo* systems were protocadherin 17 (*Pcdh17*) and semaphorin 7A (*Sema7a*), both membrane-bound structural proteins that play important roles in neuron morphology and development(53–55).

Together, these findings highlight a need to better understand the molecular and phenotypic consequences of mixtures of prevalent compounds in the human exposome. It further highlights the need to test developmental exposures on both male and females to identify shared and unique responses in relevant phenotypes. Further, these findings open new avenues of research to use more complex exposures in understanding gene-environment interactions that may be relevant to complex disorders and neuronal function.

## Materials and Methods

### Integrative Genome-Exposome Method (IGEM)

IGEM implements a configurable, stepwise pipeline for integrating external exposure-related knowledge sources and identifying exposure–exposure (ExE) interactions. Through an administrative interface, users specify data sources of interest and define which fields or columns should be mapped to exposure terms. Data ingestion is performed in discrete stages, including data retrieval, identification of exposure-related terms, and normalization to a curated internal vocabulary using predefined identifiers and aliases.

Within each record, all normalized exposure terms identified in the same context are paired to generate candidate ExE interactions. Each unique exposure–exposure interaction is counted once per record to avoid redundancy due to repeated mentions. Interactions are then aggregated across records within each data source, resulting in a single stored ExE relationship per source, annotated with the number of supporting records. Aggregated interactions are stored in a lightweight SQLite database optimized for downstream exploration, where additional interfaces allow users to filter, summarize, and query interactions of interest for analysis and visualization.

### ldEDC Solution

ldEDC solution was prepared in PBS as detailed in **Supplemental Table 1**. PFOS, DBP, and DEHP were dissolved in 100µL DMSO before slow addition to PBS for stock solution and subsequent dilution into the mixture solution. BPA and BPS were dissolved into 1mL ethanol before addition of PBS and dilution into the mixture solution. The final solution was aliquoted and kept protected from light at RT.

### Mice and ldEDC Diet Administration

All experiments were conducted in accordance with and approval of the University of Pennsylvania Institutional Animal Care & Use Committee. All mice used were on the C57BL/6J background and housed in a 12-hour light–dark cycle. Adult female mice (∼60 days) were purchased from Jackson Laboratories and housed in standard-sized cages in temperature and humidity-controlled rooms and acclimated for 2 weeks prior to being placed on custom diets. Custom diets were generated by Teklad (inotiv) by supplying the requisite weights of each chemical mixed thoroughly into solid sucrose as detailed in **Supplemental Table 1**. None of the components were dissolved into DMSO or ethanol. The vehicle and ldEDC supplements were added to AIN-93G low phytoestrogen diet with increased vitamin and mineral levels to aid in reproduction. Solid diet pellets were stored in at 4°C while not in use. Pellets in cages were replaced twice a week and appetence was monitored once a day.

Females were mated by placing two in a cage with an adult (∼100 day) C57BL/6J male mouse for 3 days. One day prior to the addition of females, the used bedding of the males was mixed into their cage to induce estrus. The diet in the male cages was temporarily replaced with the vehicle or ldEDC custom diets during mating. The F1 generation offspring were used as subjects with postnatal day 1 designated separately for each litter. Mothers and the F1 generation were kept on the custom diets until they were euthanized.

### Primary cell culture

Cortices were dissected from E16.5 C57BL/6J embryos and cultured in neurobasal medium (Gibco 21103049) supplemented with B27 (Gibco 17504044), GlutaMAX (Gibco 35050061), penicillin-streptomycin (Gibco 15140122) in 24-well plates coated with 0.05 mg/mL Poly-D-lysine (Sigma-Aldrich A-003-E). At 3 DIV, neurons were treated with 0.5 µM AraC. A total of 12 pups were sacrificed and split into 3 equal sized groups of dissociated neurons for separate seeding as replicates. At 5 DIV, half the media was replenished for each well and half the wells for each replicate were treated with 2µL either vehicle or ldEDC (a 1/1000 dilution in the 2mL media). Neurons were cultured for an additional 5 days prior to collection.

### KCl neuron activation

Neurons were cultured and exposed under the same conditions at the same timepoints as above. At 10 DIV, half the media per well was collected and 1µL of 1mM TTX (a 1/1000 dilution) and 10µL of 10mM D-AP5 (a 1/100 dilution) to quiesce the cells. 500µL of isotonic KCl solution (final concentrations: 170mM KCl, 2mM CaCl2, 1mM MgCl2, 10mM HEPES pH 7.9, in warmed, conditioned neurobasal media) was prepared and added to half of each plate. All media was removed after 30 minutes, cells were washed twice in PBS, then RNA was immediately collected.

### Bulk RNA-sequencing

#### Library preparation and sequencing

RNA was isolated using Zymo Quick-RNA Miniprep Plus Kit (R1057). 300ng per replicate, per condition were used to generate RNA-seq libraries using the Mercurius BRB-seq library preparation kit for 96 samples (Alithea Genomics 10813). Prior to sequencing, library size distribution was confirmed by capillary electrophoresis using an Agilent 4200 TapeStation with high sensitivity D1000 reagents (5067-5585), and libraries were quantified by qPCR using a KAPA Library Quantification Kit (Roche 07960140001). Libraries were sequenced on an Illumina Novaseq6000 S1 with 1% PhiX spike-in (90-bp read length, paired end).

#### Data processing and analysis

Reads were mapped to Mus musculus genome build mm10 with STAR (v2.7.1a) and assigned to exonic features using the featureCounts function of subRead (v2.0.3). DESeq2 (v1.38.0) was used for pairwise differential gene expression analysis using a negative binomial model with default model fitting parameters. 3 replicates were used for vehicle and exposure. Subsetting filters were applied to determine significant DEGs as follows: baseMean of normalized counts > 20, adjusted p value ≤ 0.05, and |fold change| ≥ 1.2 (|log2(FC)| ≥ 0. 263). Gene ontology analysis was performed using gProfiler2(56) (v0.2.3) over-representation analysis against the GO molecular function, cellular component, and biological process databases. Multiple testing was corrected using the g:SCS algorithm, a GO-optimized method that provides a better threshold between significant and non-significant than FDR(57). Only terms with ≥ genes were chosen for specificity. The domain scope of GO testing used a background list of all genes expressed in the vehicle control neurons (baseMean of normalized counts > 20).

#### Behavioral assays

Male and female vehicle treated or ldEDC exposed mice were tested as follows (n = 8 per condition, per sex). Mice were used at 4 weeks old at the onset of behavior testing which included elevated zero maze, open field, and social choice, in that order.

#### Elevated zero maze

The elevated zero apparatus consists of a circular shaped platform raised approximately 16 inches above the floor. Two opposing quadrants have raised walls (wall height = 4 inches, circle width = 2 inches) without a ceiling leaving these closed quadrants open to overhead light. The two remaining opposing quadrants were open (wall height = 0.25 inches). Mice were placed into a closed quadrant and allowed to freely explore for 5 minutes. The entire testing session was recorded, and videos were analyzed automatically using EZtrack(58) (v1.2).

#### Open field

Mice were placed into an empty arena (15 inches x 15 inches) and allowed to freely explore for 10 minutes. Videos were analyzed automatically using EZtrack, with the boundaries between periphery and open space specified by beam break ranges recorded using Photobeam Activity System Open Field software (San Diego Instruments).

#### 3-chamber social choice assay

The social choice test was carried out in a three-chambered apparatus, consisting of a center chamber and two outer chambers. Before the start of the test and in a counter-balanced manner, one end chamber was designated the social chamber, into which a stimulus mouse would be introduced, and the other end chamber was designed the nonsocial chamber. Two identical, clear Plexiglas cylinders with multiple holes to allow for air exchange were placed in each end chamber. In the habituation phase of the test, the experimental mouse freely explores the three chambers with empty cue cylinders in place for 10 min. Immediately following habituation, an age and sex-matched stimulus mouse was placed in the cylinder in the social chamber while a rock was simultaneously placed into the other cylinder in the nonsocial chamber. The experimental mouse was tracked during the 10 min habituation and 10 min social choice phases. All testing was recorded, and videos were analyzed manually by an investigator blinded to the conditions and sex of the mice.

### Single nuclei RNA-sequencing (snRNA-seq)

#### Nuclei isolation

For each biological replicate (male and female, vehicle and exposure, 4 mice each condition), one cortical hemisphere of a mouse was dissected, further microdissected to separate the prefrontal cortex from the amygdala and hippocampus, flash frozen in liquid nitrogen, stored at −80 °C. The nuclei isolation procedure used was modified from Mo (2015)(59). Tissue was homogenized in douncers using a loose pestle (∼10-15 strokes) in 1.2 mL of homogenization buffer supplemented with 1 mM DTT, 0.15 mM spermine, 0.5 mM spermidine, RNasin ® Plus Ribonuclease Inhibitor (Promega N2611), and EDTA-free protease inhibitor (Roche 11873580001). A 5% IGEPAL-630 solution was added (107µL), and the homogenate was further homogenized with the tight pestle (∼10-15 strokes). The sample was then mixed with 1.3mL of 50% iodixanol density medium (Sigma D1556) and added to a standard 15mL polypropylene tube on top of a cushion of 40% and 30% optical gradient. Tubes were spun at 4000xg on a swinging bucket centrifuge at 4°c for 20 minutes with no brake. Nuclei were then strained through a 40µm mesh into a 1.5mL tube and counted using a hemocytometer. 1 million nuclei per mouse were transferred to 15mL polypropylene tubes coated with BSA and fixed using the Evercode Nuclei Fixation kit v3 (Parse Biosciences UM0030) before being frozen at -80°c using an isopropanol-mediated cell freezing container (ThermoFisher 5100-0001).

#### SPLiT-seq library preparation and sequencing

All nuclei were thawed simultaneously and counted using a Countess 3 automated cell counter (ThermoFisher). 3200 nuclei were removed from each sample and processed using the Evercode WT v3 single nuclei isolation kit (Parse Biosciences). 8 sublibraries each with an equal representation of all samples were generated with a target nuclei count per sublibary of 6250 (50,000 total). Prior to sequencing, library size distribution was confirmed by capillary electrophoresis using an Agilent 4200 TapeStation with high sensitivity D5000 and D1000 reagents (5067-5585), and libraries were quantified by qPCR using a KAPA Library Quantification Kit (Roche 07960140001). Libraries were sequenced on an Illumina NovaseqX 10B kit with 10% PhiX spike-in (64-bp read length, paired end).

#### Data processing

Sublibraries and samples were demultiplexed using the split-pipe pipeline with default parameters which filters for nuclei with ≥ 90% confidence for each of the three barcodes. The filtered DGE output from the split-pipe pipeline (v1.5.0) aggregating all samples was converted to a Seurat object using ReadParseBio(). This was then filtered as follows: min.cells ≥ 20 per sample, min.features ≥ 200, nfeatures_RNA ≥ 200, nfeatures_RNA ≤ 40000, percent.mt ≤ 0.1, percent.ribosomal ≤ 0.5. Counts were normalized using a scale factor of 10000. Dimensionality reduction and clustering was performed using an iterative random forest method with CHOIR(60) (v0.3.0) using the first 10 principal components. The maximum number of hierarchal trees was set to 10 to minimize overclustering. CHOIR clustering was performed using an alpha of 0.05 with 100 iterations and Bonferroni correction for multiple testing. Likely doublets were identified using an artificial k nearest neighbors method via DoubletFinder(61) (v2.0) before removal. Cluster assignments were initially generated using the MapMyCells tool of the Allen Brain Atlas (RRID:SCR_024672). Marker gene expression was used to collapse like-subclusters of inhibitory neurons or non-neuronal cells. Cortical layers were refined using gene assignments in the Allen Brain Atlas.

### Downstream Analysis

#### Differential gene expression analysis

The Seurat object was subset by sex (female or male only), exposure (vehicle or ldEDC only), neuron type (excitatory or inhibitory), or non-neuron type (oligodendrocytes, astrocytes, microglia). Differential gene expression pairwise comparisons were performed using MAST(62) or DElegate(63), a pseudo-bulk methodology using EdgeR, factoring each mouse per condition as the latent variables. For MAST, DEGs were filtered for those with an adjusted p ≤ 0.05 and a minimum percentage expression per condition or cluster of 5%. For the pseudobulk, the individual mice per condition were used as latent variables to control sample variability. Only those genes that were significantly up or downregulated in both the MAST and pseudobulk methodologies were used for any downstream processing steps. Known sex-specific genes (*Xist*, *Uty*, *Kdm5d*, *Eif2s3y*, *Tsix*, *Ddx3y*, *Kdm5c*, *Eif2s3x*) were blacklisted as minor variation in the female/male ratio in a cell type or cluster would show a disproportionate change in their expression that could confound results. Gene ontology analysis was performed using gProfiler2(56) (v0.2.3) over-representation analysis against the GO molecular function, cellular component, and biological process databases as well as KEGG. STRING(64) (v12.0) was used for protein network profiling using a high confidence (0.7) threshold and only allowing the active interaction sources of experiments, databases, neighborhood, and gene fusions.

#### CellChat

CellChat(43) (v2.1.0) was used to infer and compare cell-cell communication networks across the four conditions. The full manually curated database of mouse ligand-receptor interactions (v2) was analyzed against the combined single nuclei dataset by projecting the gene expression data onto the protein-protein interaction networking using a diffusion process to reduce dropout effects of lowly-expressed ligands or receptors. Significant communication pathways were assigned to each dataset by performing a permutation test via the ‘trimean’ statistical method, approximating 25% truncated means. The identified interactions were further filtered by only including those found in groups of 10 or more cells. Information flow was calculated using rankNet(), only factoring in those pathways that contributed to ≥ 25% of the total signaling weight to filter out pathways with too few interactions. Signaling network similarities were calculated using computeNetSimilarityPairwise() based on functional similarity. Deviating pathways were separated using manifold learning.

## Funding

National Institutes of Health grant 1DP2MH129985 (EK)

National Institutes of Health grant R01NS134755 (EK)

Autism Spectrum Program of Excellence at the University of Pennsylvania

Eagles Autism Foundation

National Institute of Environmental Health Sciences T32-ES019851 (AP)

## Author Contributions

Conceptualization: AP, EK

Methodology: AP, EK, RAS, YCL, ALGR, NP, MH

Investigation: AP, RA, CQ, ALGR, NP

Visualization: AP

Formal Analysis: AP

Supervision: EK

Writing – Original Draft: AP

Writing – Review and Editing: AP, EK, RAS, YCL

## Data and materials availability

RNA-sequencing and single nucleus RNA-sequencing data generated in this study will be available under the following GEO accession numbers GSE298626 and GSE298627 upon publication. All data are available in the main text or the supplementary materials. Any additional data will be made available within two weeks upon request to corresponding author.

## Statistical analysis

All statistical analysis were performed using readily available code in R. The number of replicates and details of statistical tests are reported in figure legends and methods. Shapiro-Wilk’s test was used to determine the normality of a given dataset.

---

**Supplemental Figure 1:**
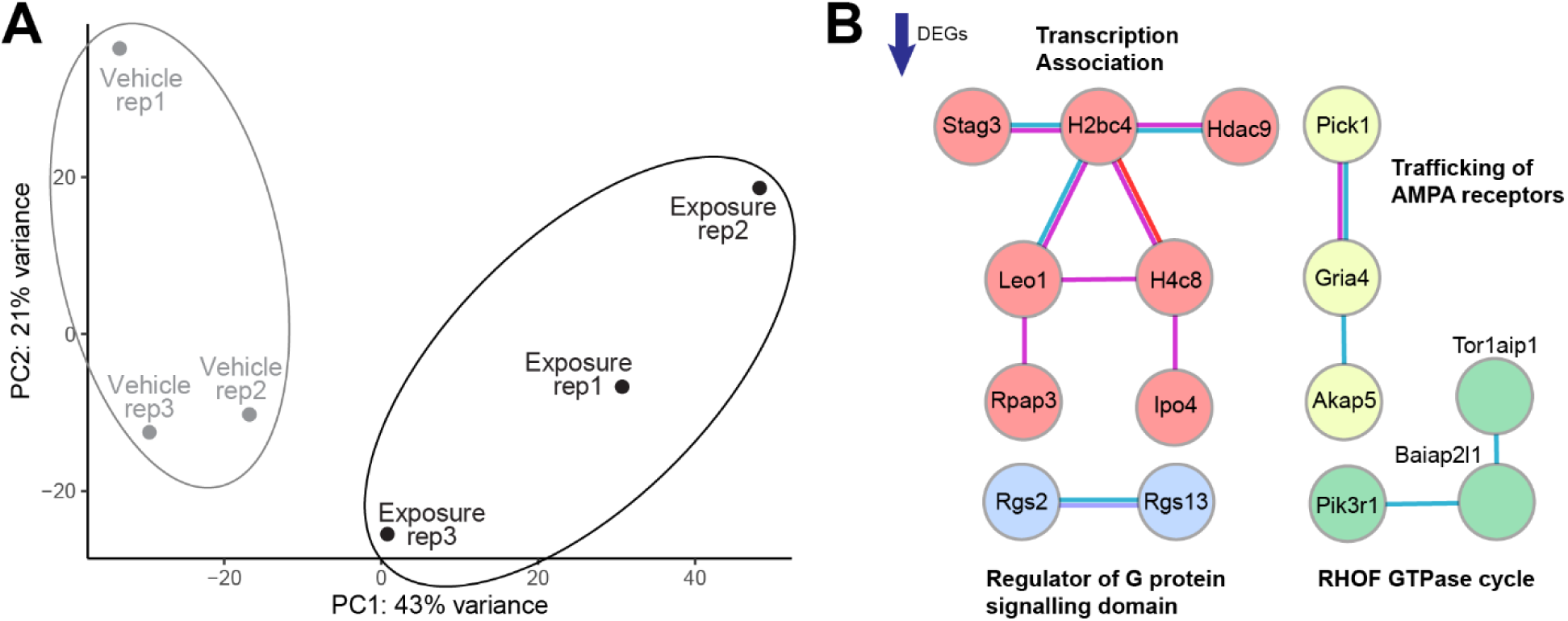
Additional analysis of neuron ldEDC exposure RNA-seq. **A**. Principal component analysis of the pairwise comparison between vehicle and ldEDC-treated neurons (n = 3). **B**. STRING analysis of significantly downregulated genes in the ldEDC treatment. Only genes with at least one interaction passing a high confidence score (0.7) are shown. Modules were determined by MCL clustering with an inflation parameter of 3. Edge colors represent different interaction categories (purple: experimental, blue: databases, green: protein neighborhood, red: protein fusions, dark blue: co-occurrence).

**Supplemental Figure 2:**
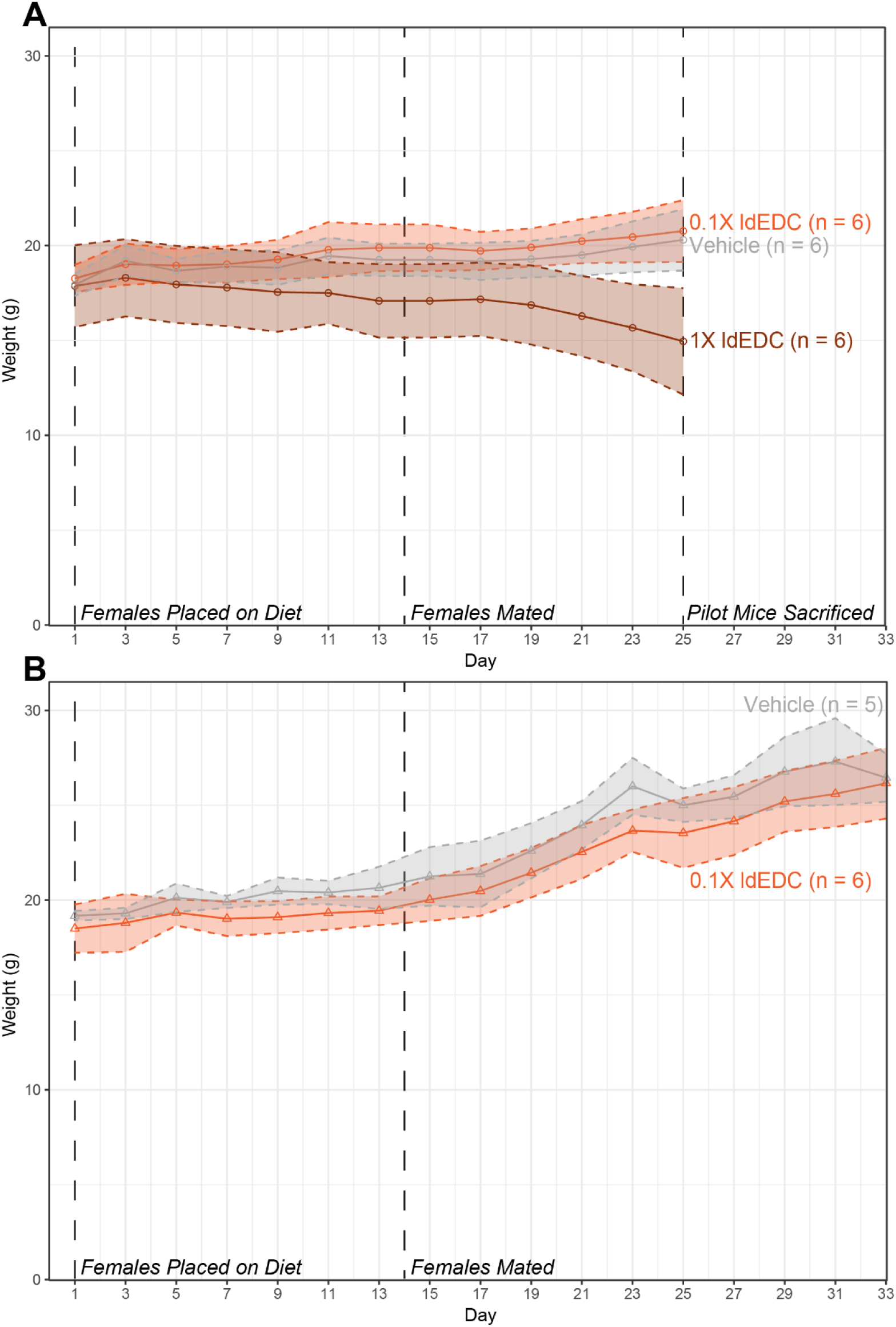
Exposed female weights during diet administration. **A**. Weight (in grams) of female mice after being placed on low-phytoestrogen diet with 0X, 0.1X, or 1X ldEDC in the pilot experiment (n = 6 for each diet). **B.** Weight (in grams) of female mice that generated the F1 generation for downstream testing after being placed on low-phytoestrogen diet with 0X or 0.1X ldEDC (vehicle: n = 5, 0.1X exposure: n = 6).

**Supplemental Figure 3:**
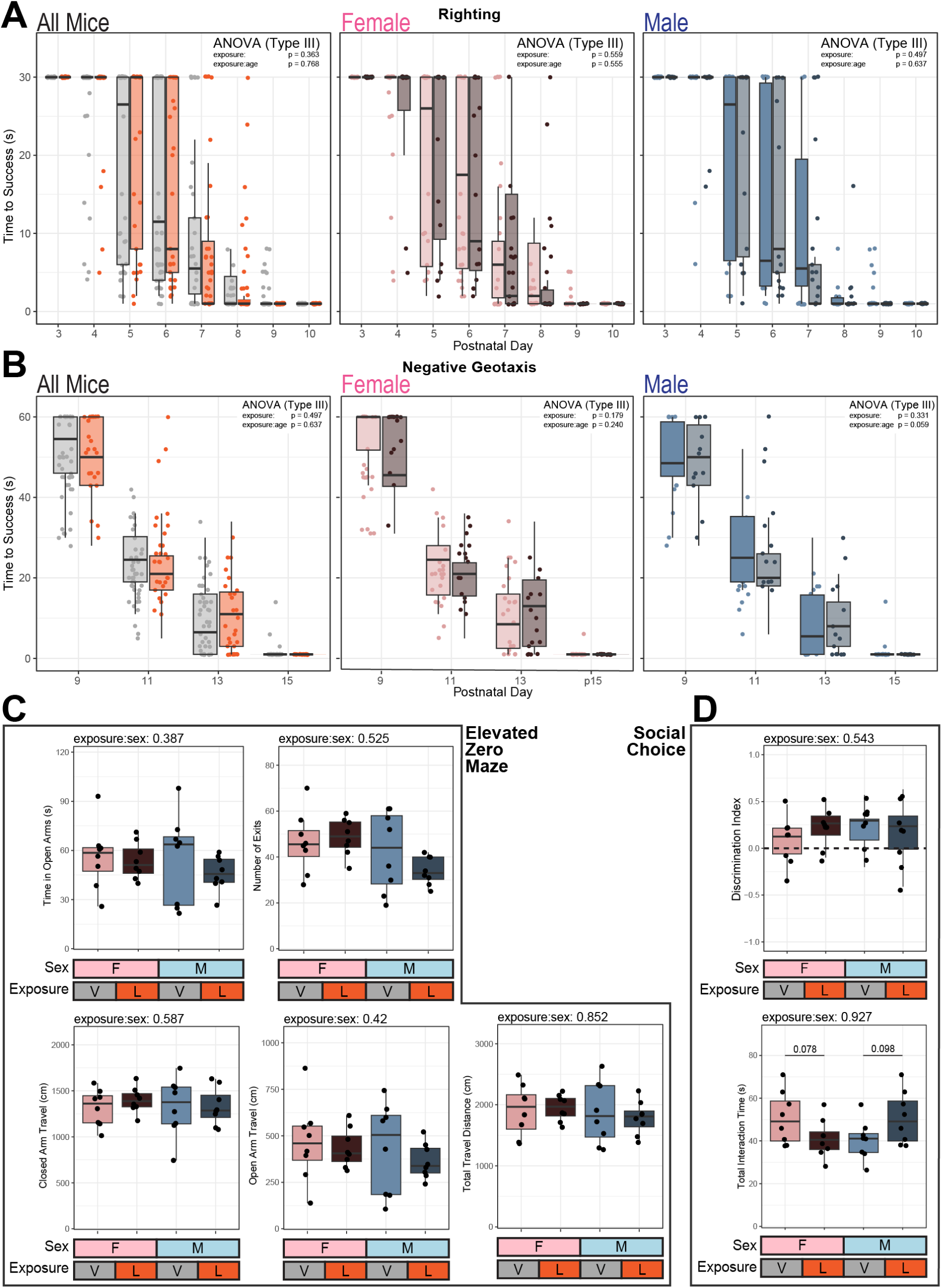
Additional development and behavior test of perinatal vehicle or ldEDC exposed mice. **A**. Time to surface righting in male and female pups by exposure (male[vehicle: n = 21, exposure: n = 14]; female[vehicle: n = 22, exposure: n = 16]). Type III ANOVA results shown above results with all mice or separated by sex. Time to success was determined as how long it took for a pup turned onto its back on a flat surface to turn over onto its belly. If the pup took ≥ 30 seconds, their time was recorded as 30. **B**. Time to right self in negative geotaxis assay in male and female pups by exposure. Time to success was determined by how long it took for a pup to change its orientation up an inclined plane when placed facing downward. **C**. Measures from the elevated zero maze (repeated measures ANOVA above each). **D**. Measures from the social choice assay (repeated measures ANOVA above each, Kruskal-Wallis Test between pairs).

**Supplemental Figure 4:**
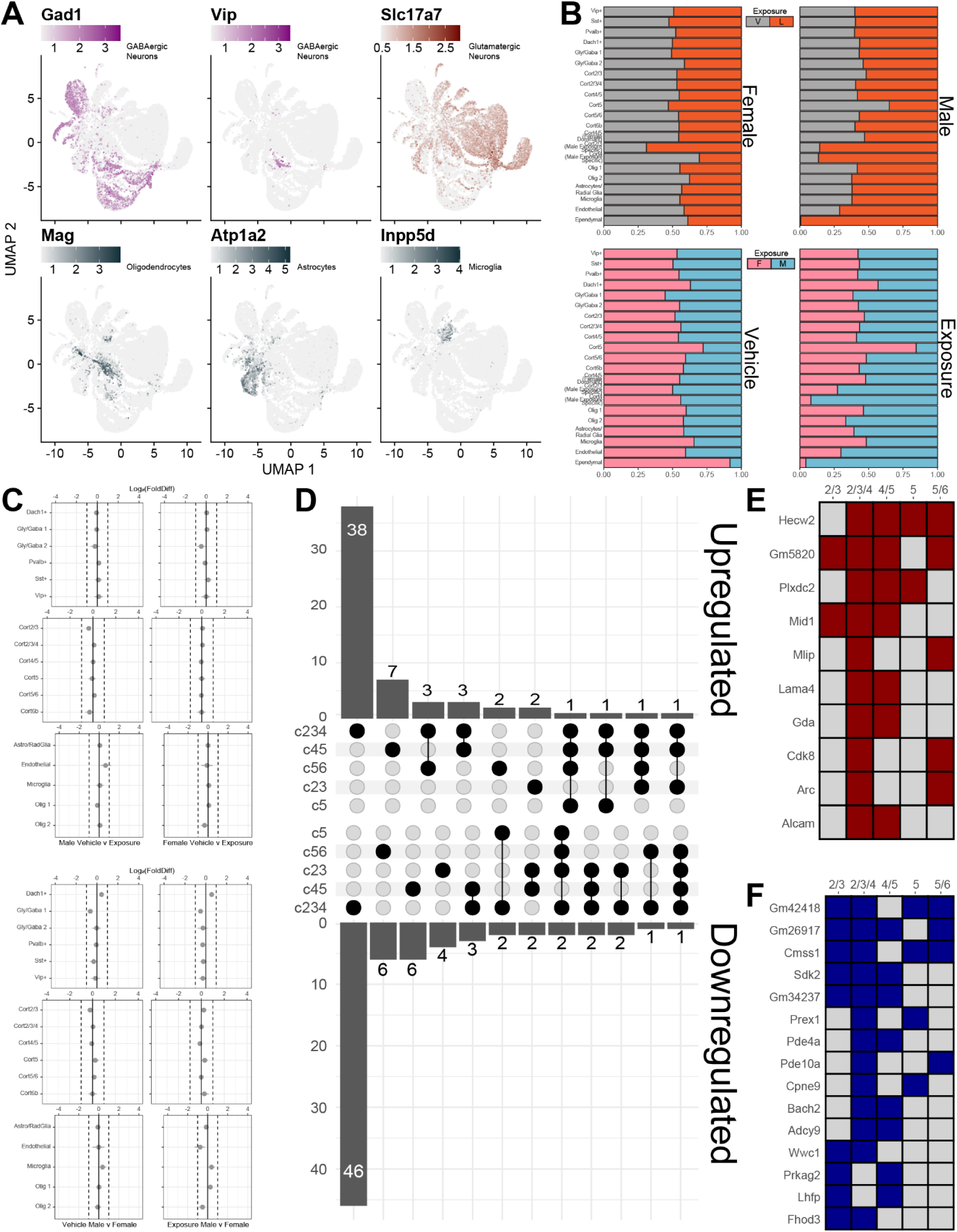
Cluster-specific comparisons. **A**. Featureplots of marker genes for GABAergic neurons (*Gad1*, *Vip*), glutamatergic neurons (*Slc17a7*), oligodendrocytes (*Mag*), astrocytes (*Atp1a2*), and microglia (*Inpp5d*). **B**. Stacked bar charts of the proportion of each exposure condition for each cluster separated by sex (top) or each sex separated by exposure (bottom). **C**. Proportion of nuclei in each cluster comparing vehicle and exposure nuclei separated by sex (top) or male and female separated by exposure (bottom). Grey indicates false discovery rate ≥ 0.05. **D**. UpSet plot of the number of differentially up or downregulated genes found in one or more excitatory cell cluster. **E-F**. DEGs up (**E**) or downregulated (**F**) in at least 2 excitatory cell clusters.

**Supplemental Figure 5:**
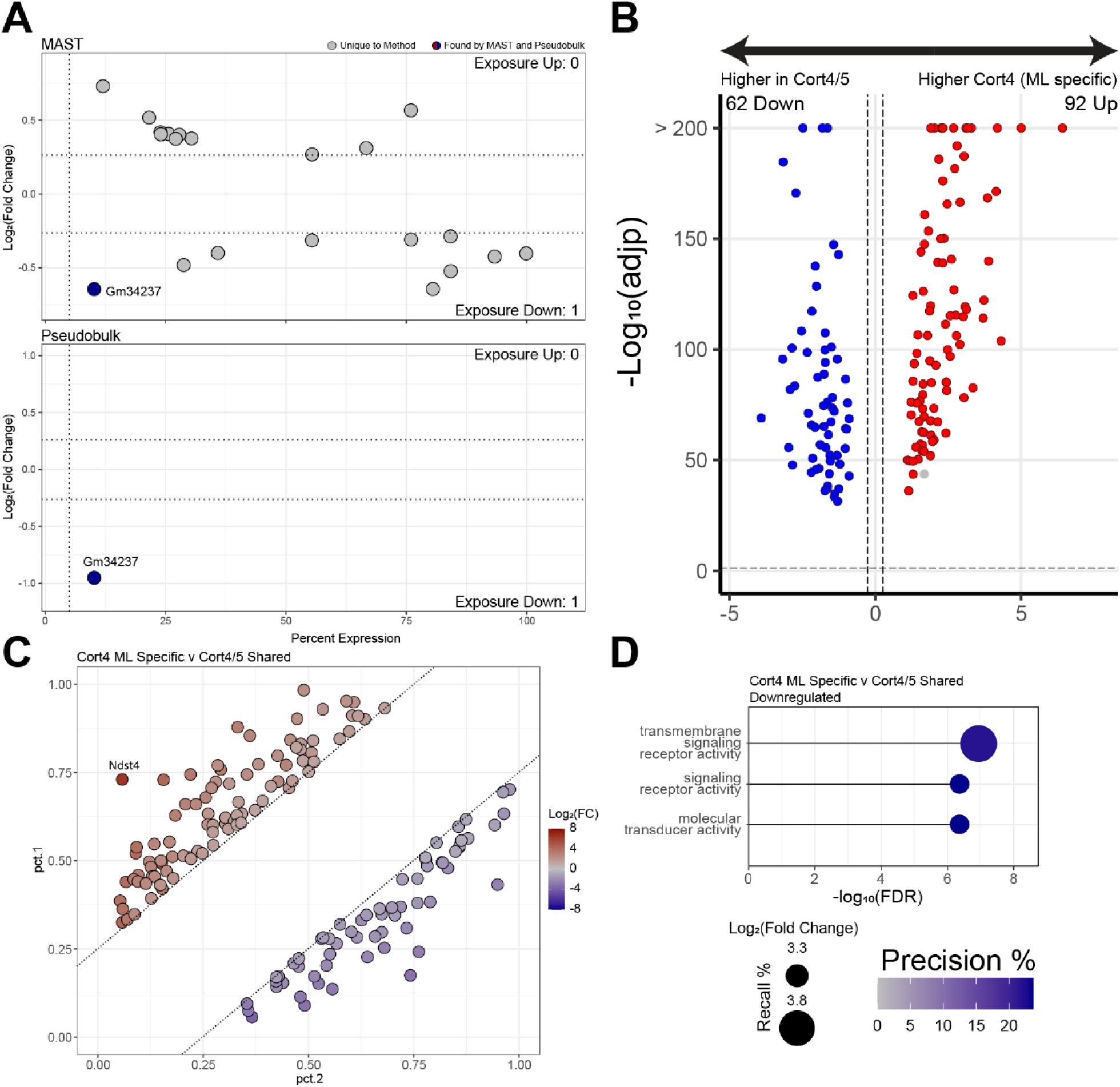
Additional analysis of the single nuclei dataset. **A.** Scatter plots of differentially expressed genes between exposure and vehicle for all inhibitory neurons. Points are colored if the gene was significant (|fold change| ≥ 1.2, adj(p) ≤ 0.05) in both the MAST and pseudobulk analysis. **B**. Volcano plot of genes differentially expressed in the ML-specific cortical 4 excitatory neuron cluster compared to the condition-balanced cortical 4/5 cluster. **C**. Scatterplot of genes by their percent expression in the ML-specific cortical 4 cluster (pct.1) or condition-balanced cortical 4/5 cluster (pct.2). Only those genes with an |fold change| ≥ 1.2, adj(p) ≤ 0.05, and minimum pct = 0.25 are shown. **D**. Gene ontology analysis of significantly downregulated differentially expressed genes between the cortical 4 clusters. Recall is the proportion of functionally annotated genes in the query over the number of genes in the GO term. Precision is the number of genes found in the GO term over the total number of genes in the query.

**Supplemental Figure 6:**
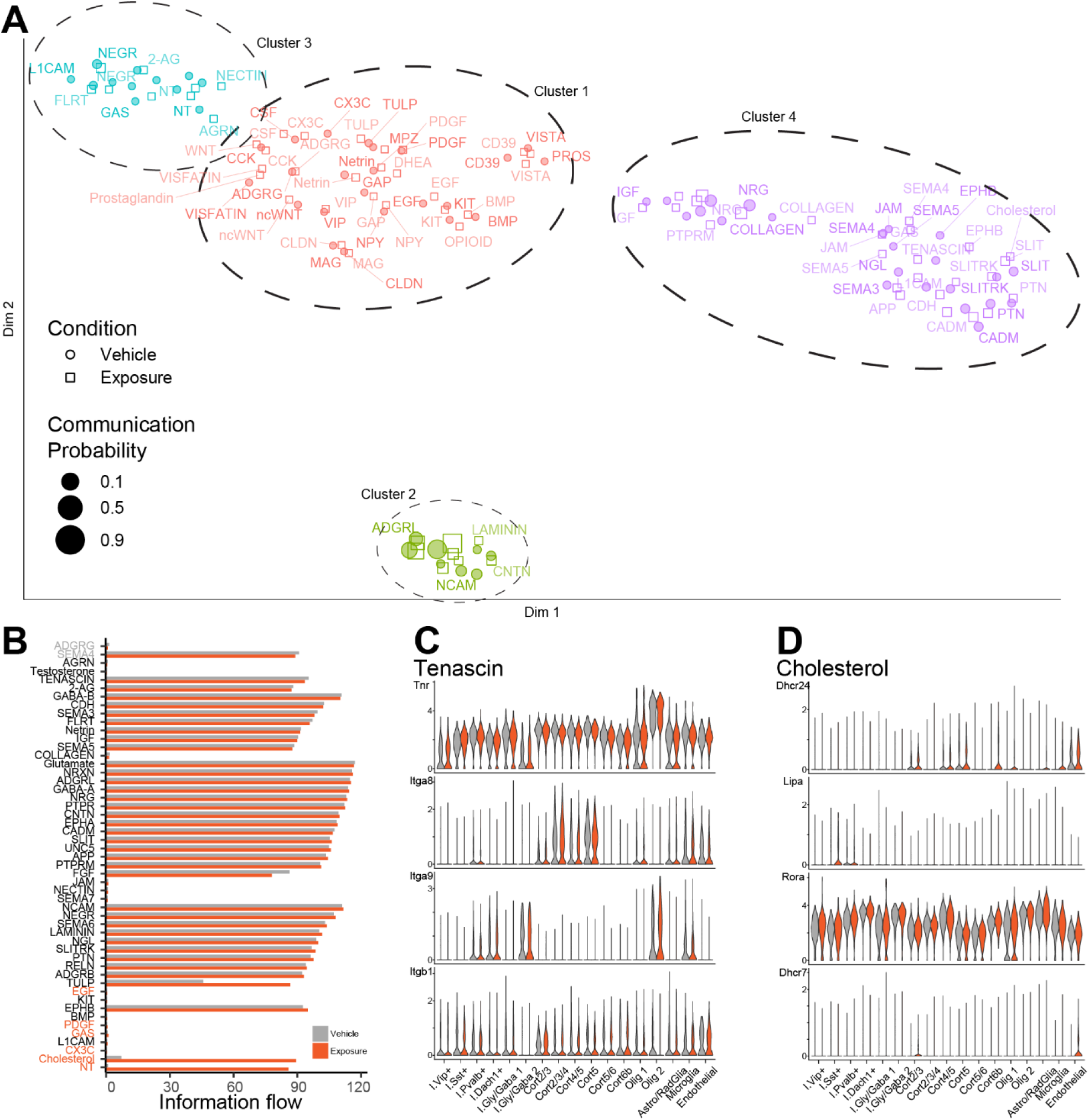
CellChat metrics. **A**. 2-D visualization of signaling pathways that significantly differ between vehicle and exposure conditions. Pathways are clustered by functional similarity based on major senders and receivers. **B**. Bar plot of the total information flow from signaling pathways contributing to ≥ 25% of the total signaling weight found in vehicle or exposure conditions. Colored pathway names indicate those significantly enriched for one condition over the other. **C-D**. Violin plots of normalized gene expression for components of the tenascin (**C**) and cholesterol (**D**) signaling pathways by cluster.

**Supplemental Figure 7:**
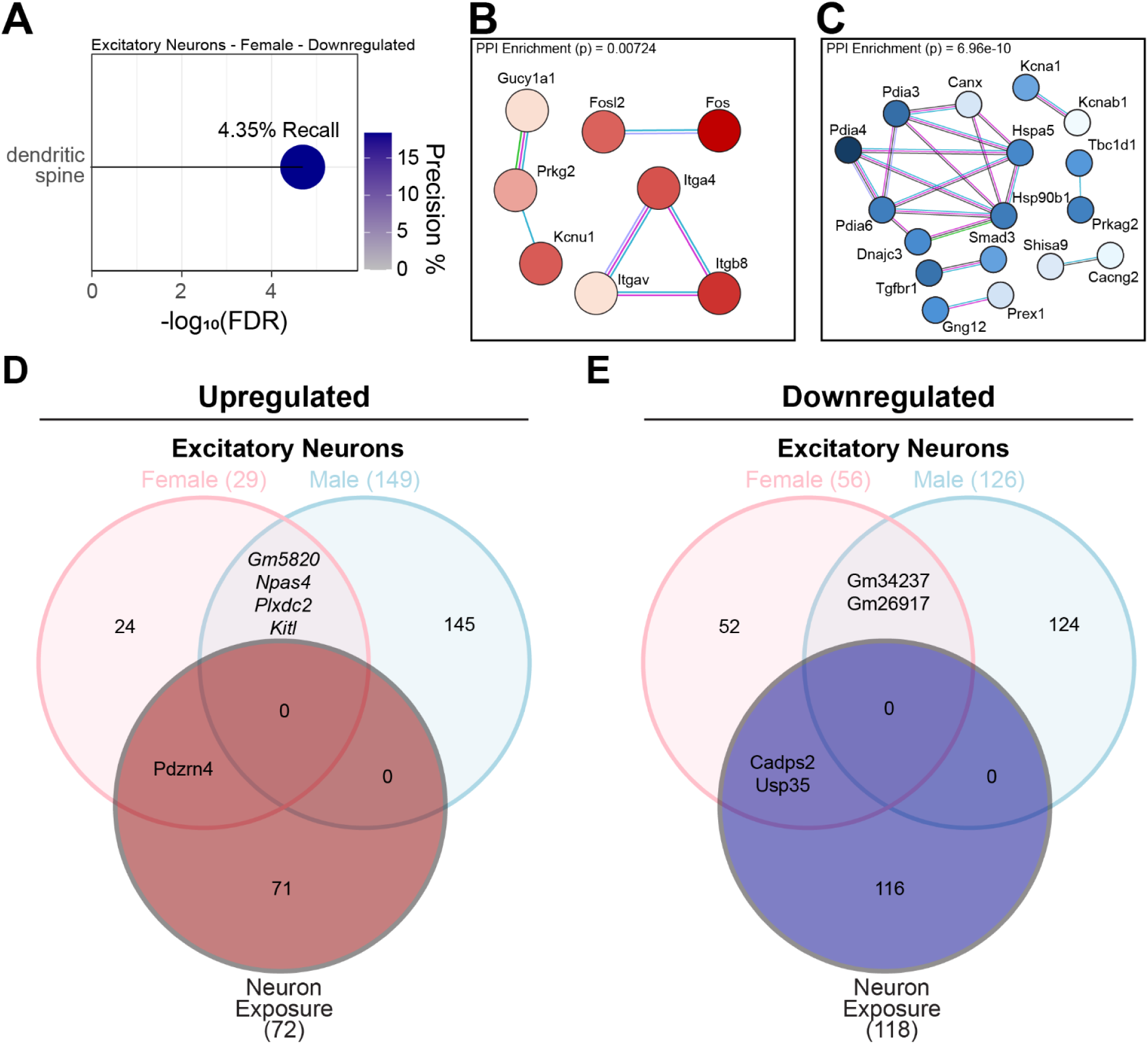
Sex-specific DEG profiling. **A**. Gene ontology analysis of female-specific downregulated DEGs in excitatory neurons. Recall is the proportion of functionally annotated genes in the query over the number of genes in the GO term. Precision is the number of genes found in the GO term over the total number of genes in the query. **B-C**. Significant STRING protein-protein interaction modules among male-specific up (**B**) and downregulated (**C**) DEGs. Nodes are colored by relative log_2_FC. Edge colors represent different interaction categories (purple: experimental, blue: databases, green: protein neighborhood, red: protein fusions, dark blue: co-occurrence). **D-E**. Overlap of genes up (**D**) or downregulated (**E**) in cultured neurons following ldEDC exposure as in Figure 1 against excitatory neurons stratified by sex.

**Supplemental Data Table 1. ldEDC solution and diet recipes**

**Supplemental Data Table 2. Neuron bulk RNA-seq differentially expressed genes**

**Supplemental Data Table 3. RNA-seq gene ontology and motif analyses**

**Supplemental Data Table 4. Cortical single nuclei RNA-seq differentially expressed genes by cluster and analysis type**

**Supplemental Data Table 5. Male-exposure-specific cluster marker genes**

**Supplemental Data Table 6. Single nuclei RNA-seq gene ontology analysis**

**Supplemental Data Table 7. CellChat pathway scoring by condition**

**Supplemental Data Table 8. Cortical single nuclei RNA-seq differentially expressed genes by sex, cell type, and analysis type**

